# Single-cell network biology characterizes cell type gene regulation for drug repurposing and phenotype prediction in Alzheimer’s disease

**DOI:** 10.1101/2022.01.09.475548

**Authors:** Chirag Gupta, Jielin Xu, Ting Jin, Saniya Khullar, Xiaoyu Liu, Sayali Alatkar, Feixiong Cheng, Daifeng Wang

## Abstract

Dysregulation of gene expression in Alzheimer’s disease (AD) remains elusive, especially at the cell type level. Gene regulatory network, a key molecular mechanism linking transcription factors (TFs) and regulatory elements to govern target gene expression, can change across cell types in the human brain and thus serve as a model for studying gene dysregulation in AD. However, it is still challenging to understand how cell type networks work abnormally under AD. To address this, we integrated single-cell multi-omics data and predicted the gene regulatory networks in AD and control for four major cell types, excitatory and inhibitory neurons, microglia and oligodendrocytes. Importantly, we applied network biology approaches to analyze the changes of network characteristics across these cell types, and between AD and control. For instance, many hub TFs target different genes between AD and control (rewiring). Also, these networks show strong hierarchical structures in which top TFs (master regulators) are largely common across cell types, whereas different TFs operate at the middle levels in some cell types (e.g., microglia). The regulatory logics of enriched network motifs (e.g., feed-forward loops) further uncover cell-type-specific TF-TF cooperativities in gene regulation. The cell type networks are highly modular. Several network modules with cell-type-specific expression changes in AD pathology are enriched with AD-risk genes and putative targets of approved and pending AD drugs, suggesting possible cell-type genomic medicine in AD. Finally, using the cell type gene regulatory networks, we developed machine learning models to classify and prioritize additional AD genes. We found that top prioritized genes predict clinical phenotypes (e.g., cognitive impairment). Overall, this single-cell network biology analysis provides a comprehensive map linking genes, regulatory networks, cell types and drug targets and reveals mechanisms on cell-type gene dyregulation in AD.

## Introduction

Alzheimer’s Disease (AD) is a brain disorder that progresses into memory loss, a decline in cognitive skills, and ultimately dementia. The mechanistic causes of AD are not yet fully understood, especially at the cell type level, although the abnormal accumulation of neuronal tangles and amyloid plaques in the AD brain have become potential hallmarks of the disease. The genetic factors that possibly lie upstream of various AD phenotypes have now been extensively studied from next generation sequencing data, such as genome-wide gene expression changes. A variety of computational analyses have been applied to those data for understanding abnormal gene expression and regulation in AD. However, most studies have been performed on bulk tissue data and missed cell-type-specific signals. The neurovascular unit as a whole could drive AD progression^1^, and recent studies have verified that molecular changes in AD are highly cell-type-specific^2^. Thus, it is imperative to investigate the contribution of individual cell types in the brain to the progression of AD along with clinical phenotypes. Emerging single-cell RNA-seq enables such an analysis, as it captures the transcriptomic landscape of individual cells, offering a rich source of data for the analysis of dysregulated molecular systems within individual cells.

Several studies have highlighted strong links between molecular connectivity and human diseases, suggesting that disease risk genes often work together as a coherent biological network. Thus, it is critical to study broken functional relationships between genes, rather than individual genes, to better understand the molecular mechanisms associated with the disease. Network biology offers a powerful computational framework that transcends individual gene investigation that uses univariate methods, such as differential expression analysis. For example, gene regulatory networks (GRNs) provide information about regulatory interactions between regulators, e.g., transcription factors (TFs), and their potential target genes. Such GRN models can be used to derive novel biological hypotheses about dysregulated disease pathways. With scRNA-seq data quickly accumulating in open repositories, single-cell network biology is now leading a shift from the traditional bulk RNA-seq mediated analyses^3–7^. Although GRNs in AD have been previously explored using expression data from bulk tissues^8–13^, cell type level GRN in AD remains under-investigated, especially via network biology approaches.

Network biology has been successfully applied to prioritize novel disease genes. The basic idea is to identify regulatory genes that have more influence over the network by virtue of their network position. Naturally, a more prominent position in the network is occupied by hubs or genes with a relatively larger number of connections and those that facilitate signaling between distant genes in the network. Hubs play a central role in modulating the expression of many genes and thus biological processes and pathways. Network biologists have adopted various classical metrics from graph theory to identify hubs in GRNs. Networkbased indicators of gene importance have also been useful for the analysis of disease at the cell type level. For example, Iacono et al. analyzed healthy and diabetic pancreatic cell networks and found that genes involved in type-2 diabetes differ in their centralities scores^14^. In addition, single-cell network analyses have revealed genes that rewire with exposure to differentiation cues^15^ and cancer-causing perturbations^16^. In the context of brain diseases, single-cell gene networks have indicated a potential cell type preference of neuropsychiatric and neurodegenerative disorders^17^ and neurodevelopmental disorders (NDDs)^18^. The authors in the later study estimated coexpression between sets of known NDD-risk genes and demonstrated that most genesets have higher coexpression in neural progenitor cells, suggesting a convergent role of these cell types in NDDs^18^.

The structure of gene regulatory networks can also help understand coordinated gene regulation. Recurring sub-graphs, called network motifs, are patterns that appear in real networks more often than random networks. Network motifs are the building blocks of biological networks. Therefore, comparing network motifs in, for example, control and disease states of the transcriptome can unravel how the disease affects the structural design of the GRN. For example, the feed-forward loop (FFL) is a three-node motif particularly interesting for analysis directed networks^19,20^. FFLs comprise a master regulator, which regulates an intermediate TF, and both TFs directly regulate the expression of a common target gene. This information can help gauge changes in ‘regulatory pressure’ on downstream TFs through coordinated activities between upstream TFs. Network motifs can also be helpful in applying digital computing ideas such as logic gates in synthetic biology^21,22^.

Network medicine is an upcoming field to solve the problem of drug repurposing by finding new uses of existing drugs by linking them to drug targets which are also implicated in human diseases^23–26^. Network-medicine approaches have been applied to repurpose drug candidates for cancers ^27^, tuberculosis^28^, and, more recently, respiratory illnesses like COVID-19^29,30^. We have also previously developed network-medicine strategies for AD. For example, we recently proposed an endophenotype network-based drug repurposing framework for AD. ^31^. Our approach uses disease-associated modules (modules enriched with disease genes) and network proximity analysis for in silico drug repurposing. Using this approach, we discovered sildenafil as a new candidate drug for AD, tested it using insurance record data, and validated it using iPSCs from patients with AD^31^. Our study shows that quantifying the network distance between AD modules and drug targets in the human interactome can significantly improve in silico drug discovery.

In this study, we developed a single-cell network biology framework to unravel the characteristics of GRNs in normal and AD conditions, discover hubs and modules of coregulated that potentially contribute to AD pathogenesis, identify disease module-associated targets of approved AD drugs, and generate a genome-wide ranking of genes according to their potential association to AD (Fig. 1). We integrated available single-cell gene expression, chromatin interaction, and TF-binding site information to infer GRNs for neuronal and glial cell types of the human brain in control and AD conditions (Fig. 1A). These GRNs depict interactions between TFs and target genes based on the level of coexpression in individual cell types and the evidence of the TF binding to the promoter or enhancer region of the target genes in open chromatin regions. We used various measures of centrality and hierarchy to outline shared and common hubs of cell type GRNs, as well as network motifs and logic gates to understand coordinated TF activities (Fig. 1B). We analyzed the modular organization of cell type GRNs and adopted a network-proximity strategy to link modules of co-regulated, biological processes, and drugs approved for AD (Fig. 1C). Finally, we utilized known AD genes from the published literature in a machine-learning strategy to generate a genome-wide ranking of genes according to their potential association to AD (Fig. 1D).

**Figure 1:**
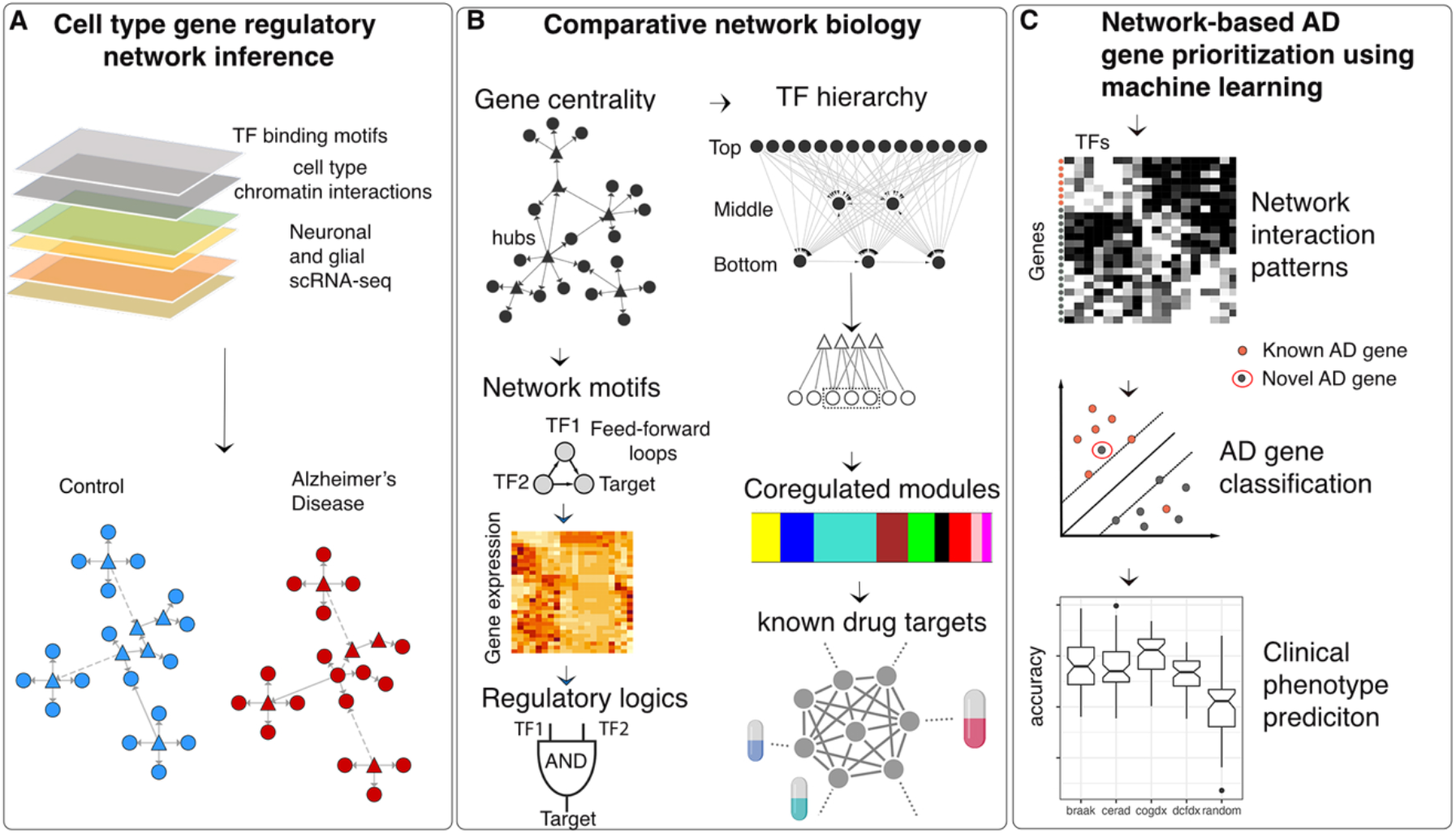
An integrative network-biology approach for analyzing cell type gene regulatory mechanisms in Alzheimer’s disease (AD). A) Predicting neuronal and glial cell type GRNs from multi-omics data in AD and control by integrating scRNA-seq with chromatin interaction data and TF binding site information. B) Analyses of cell type GRN characteristics include identification of hub genes, regulatory hierarchy, network motifs, regulatory logics, and modules of co-regulated genes. C) Machine learning based prioritization of novel AD genes using network interaction patterns and prediction of clinical phenotypes using machine learning. D) Discovery of repurposed drug target genes from cell type GRNs integrated with electronic health records.

## Results

We applied our framework to single-cell gene expression data for four cell types, including excitatory and inhibitory neurons, microglia, and oligodendrocytes from human brains diagnosed with Alzheimer’s disease (AD) and healthy controls^2^. All detailed descriptions on datasets and data processing are available in Methods. The dataset consists of single-nucleus RNA-sequencing (snRNA-seq) samples from the prefrontal cortex of 24 individuals diagnosed with AD and 24 age-matched controls with no AD pathology. In addition, we also obtained cell-type chromatin interactions^32^ and human transcription factor binding site information^33^. We integrated these datasets to infer gene regulatory networks for the four cell types in control and AD conditions. In addition to this, we also obtained single-cell transcriptomic data of healthy cells from an independent study^34^ to check the reproducibility of our results.

### Hubs of the brain cell type gene regulatory networks

We linked transcription factors (TFs), non-coding regulatory elements, and target genes to infer cell type GRNs in control and AD. Because GRNs typically have a nonuniform distribution of links (edges)^35^, system biologists are often interested in identifying ‘hub’ nodes (genes) for practical applications^36^. Hubs represent highly connected genes that have a greater influence over the network. Such highly connected hub genes often play a crucial role in modulating gene expression changes, and thus disease-associated pathways. Given that gene expression phenotype in AD is highly cell type specific^2^, we asked if distinct or similar sets of genes act as hubs across cell type networks.

We used three standard centrality metrics to quantify the influence of a given TF over each cell type’s control and AD network. The out-degree centrality calculates the number of targets for each TF, in-degree indicates how strongly a TF is under the regulatory influence of other TFs, and the betweenness centrality of a TF is a function of its out-degree and in-degree and estimates its ability to act as a communication channel between upstream regulators and downstream pathway genes. We observed that TFs with the highest out-degrees (top 10% of the sorted list) are largely common (173 TFs) across all cell types (Fig. 2A; Data S1). However, distinct TFs represent betweenness centralities of different cell type GRNs. Moreover, the overlap of such TFs is relatively higher between neuronal than glial cell types (Fig. 2B). We noted that TFs that have the greatest regulatory influence of other TFs (high incoming degrees) also vary across cell types (Fig. S1A). The overlap of such TFs is also larger between neuronal than glial cell types (Fig. S1A).

**Figure 2.**
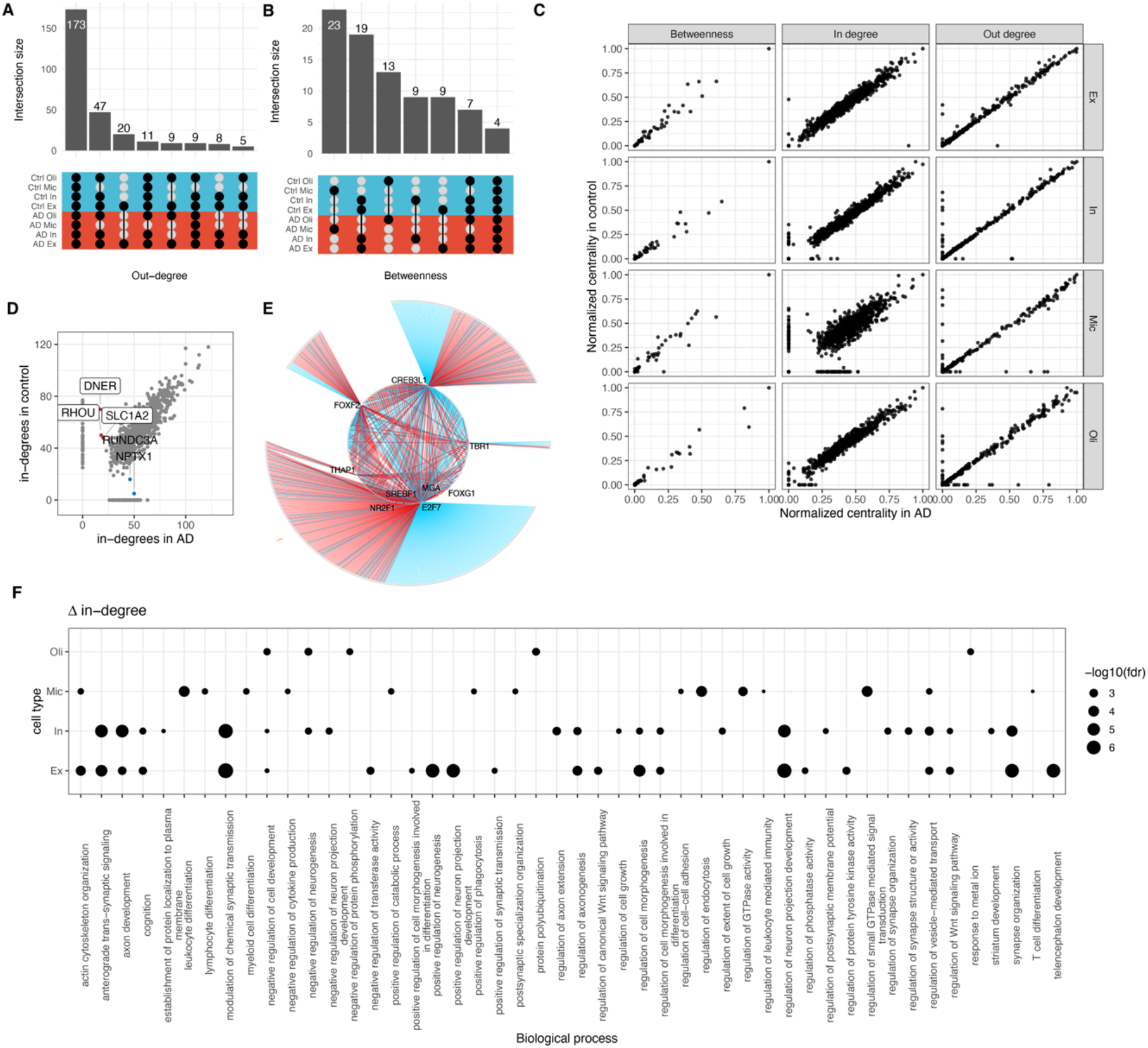
Centrality analysis reveal hub gene changes of cell-type gene regulatory networks in AD. A) An upset plot showing overlaps between the top 10% genes with the largest out-degree and B) betweenness centralities. The filled dots in the center matrix indicate the comparison between the respective sets (along the x-axis), and the bars on the top show size of the intersection. Blue and red rows indicate control and AD, respectively. C) Scatter plots showing normalized gene centralities distribution and D) the distribution of in-degrees in microglial AD and control networks. Genes with large changes in in-degree between AD and control are labelled. E) A dot plot showing enrichment of gene ontology biological processes (y-axis) among genes with the most extensive changes in the in-degree centrality across all cell types (x-axis). The dot size is set according to the FDR-corrected *p* values, as shown in the key. F) Visualization of the subnetwork of 9 TFs with high betweenness centrality in excitatory neurons. Grey circles around the periphery of the network indicate target genes. Symbols of the nine central TFs are shown and the rest hidden for clarity. Blue and red edges indicate interaction in the control and AD networks, respectively.

Although the normalized gene centralities (including non-TF genes) between control and AD GRNs across all cell types are largely correlated, there is a clear differential in the in-degrees, with the most prominent scatter in microglia (Fig. 2C). For example, the DNER, RHOU, and SLC1A2 genes have fewer regulators in AD compared to control, whereas RUNDC3A and NPTX1 are regulated by more TFs in AD than in control (Fig. 2D). DNER activates the NOTCH1 pathway which is linked to AD^37,38^. SLC1A2 mediates cellular uptake of glutamate, and loss of function of glutamate transporters has been linked to AD^39^. NPTX1 is a member of the pentraxin family, known to modulate synaptic transmission in normal conditions^40^. Also, we noted that microglia GRNs have the largest number of distinct high betweenness TFs. Therefore, we were interested to investigate if the subnetworks around these central TFs have identical or disjoint node- and edge-sets. Visualizing the network neighborhoods of the 23 high betweenness microglia TFs, we observed a considerable difference between the control and AD networks (Fig. 2E). This indicated presence of a disease-driven regulatory apparatus governed by the same TFs.

Overall, genes with the largest in-degree changes are significantly enriched in (1% FDR based on hypergeometric tests) in gene ontology (GO) biological process (BP) terms related to the immune system such as ‘neutrophil-mediated immunity’ and ‘leukocyte migration’ in microglia, development-related processes such as ‘autonomic nervous system development’ and ‘anterior-posterior pattern specification’ in oligodendrocytes, and synapse related processes such as ‘synapse organization’, ‘regulation of synaptic transmission’, and ‘modulation of chemical synaptic transmission’ in neuronal cell types (Fig. 2F).

We also checked if our centrality analysis is reproducible in an independent dataset. To do this, we obtained the dataset generated by an earlier study that analyzed gene expression in healthy brain cell types as our secondary dataset^34^. We applied our GRN inference pipeline and centrality analysis to microglia and oligodendrocytes in the secondary dataset and found that the top central genes (top 20% across all centralities) show high and statistically significant overlaps (permutation-based *P*-value = 0.00001) with the main dataset (Fig. S1B-G). Because the secondary dataset has samples only from healthy brain cells, we could only compare the overlap with our control networks. Nevertheless, the high overlaps indicated that the top central genes we identified are indeed independent of the underlying dataset and thus the approach is generalizable.

### The regulatory hierarchy of brain cell type gene regulatory networks

Given that GRNs are typically hierarchical in structure^41–43^, we asked if AD induces changes to the regulatory hierarchy of cell type GRNs. We wanted to identify TFs that act as master regulators, and other TFs that function downstream of the master regulators. Master regulators are defined as TFs at the top of the network hierarchy with no regulatory influence from other TFs^44^.

To classify TFs at different levels of regulatory hierarchy, we used the standard hierarchy height (*hh*) metric^43^. According to the *hh* metric, TFs at the top levels of the hierarchy exhibit many outgoing edges but no incoming edges (master regulators not regulated by other TF), TFs at the middle levels exhibit both incoming and outgoing edges (regulators and regulated by other TFs) and TFs at the bottom levels exhibit no outgoing edges to other TFs (highly regulated by other TFs). We found the distribution of normalized *hh* to be trimodal across all GRNs and significantly different from random networks (estimated using the KS test of 1000 random networks) (Fig. 3A, 3B, and S2A; Data S2), indicating that the brain cell type GRNs are indeed hierarchical. We also noted that the *hh* of TFs is not significantly different in control and AD networks (Fig. S2B). We found 85 (27.6%) master regulators common across all cell types AD GRNs, with the most unique master regulators in excitatory neurons (37 TFs; 12%) (Fig. S2C). Some common master regulators include known AD genes, such as CREB1, ESR1, HSF1, PPARG, NFE2L2, SPI1, TCF3, TCF7L2, TP53, CLOCK, and GLIS3. However, we found very few TFs in the middle-level (5 TFs in microglia, 3 in excitatory neurons, 1 in inhibitory neurons, and none in oligodendrocytes; Fig. S2D; see Discussion).

**Figure 3.**
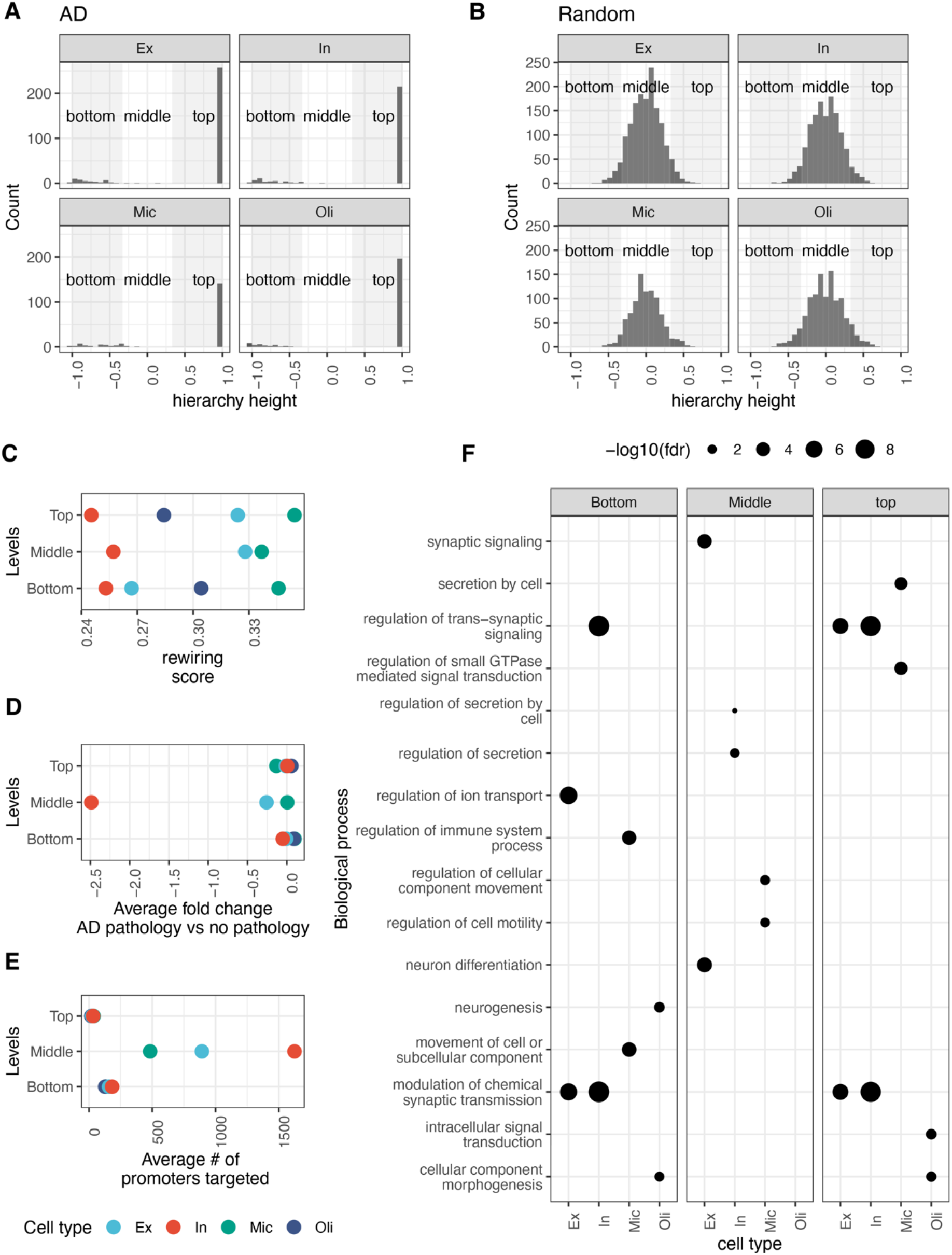
Hierarchy analysis of cell type gene regulatory networks in AD. A) The distribution of hierarchy height metric (x-axis) of TFs in AD GRNs across all four cell types. B) The distribution of hierarchy height metric (x-axis) of TFs in random networks created by preserving node-degree statistics of the corresponding AD network. C-E) Dot plots showing the rewiring scores of TFs (x-axis) across all three levels of hierarchies (y-axis), average fold change of TFs in individuals diagnosed with AD versus healthy controls, and the average number of promoters targeted by TFs at the three hierarchy levels, respectively. F) A dot plot showing enrichment of gene ontology biological processes (y-axis) within targets of top, middle and bottom layers of the regulatory hierarchy across cell type networks (x-axis).

We were interested in knowing if the readjustment of the targets of TFs at various levels of the regulatory hierarchy contributes to AD. We estimated the rewiring score of TFs based on the overlap between their predicted targets in control and AD networks (see Methods). Within the four cell types we analyzed, we found that TFs are least rewired in inhibitory neurons and most in microglia (Fig. 3C). Furthermore, on examining the differential expression in AD pathology versus no pathology (healthy controls), we found that TFs at all levels of the regulatory hierarchy generally remain stable (Fig. 3D). One down-regulated outlier is the early growth response 1 (EGR1) found in the middle-level of the inhibitory neurons. EGR1 is a known mediator and regulator of synaptic plasticity and neuronal activity and implicated in various neuropsychiatric disorders^45^. Given that the upstream regulators of EGR1 are also well characterized^46^, it is not surprising to find EGR1 in the middle-level of the inhibitory neuron’s regulatory hierarchy.

Next, we investigated the size of TF regulons in the full AD GRNs (including non-TF genes) and found that the master regulators generally target a relatively smaller number of promoters and enhancers (Fig. 3E and S3). Interestingly, middle-level TFs target many promoters (Fig. 3E). To draw a biological interpretation of the regulatory hierarchy, we performed enrichment analysis of the most confidently predicted targets of TFs using functional annotations biological process category of the human gene ontology. Interestingly, we found the top-level master regulators and the bottom-level TFs in the neuronal cell type GRNs functionally converge to regulate trans-synaptic signaling in neuronal cell types and cellular component morphogenesis in oligodendrocytes (Fig. 3F; Data S2). Master regulators in microglia seem to regulate small GTPase mediated signal transduction and secretion. We found that the middle-level TFs target distinct processes; synaptic signaling and neuron differentiation in excitatory neurons, secretion in inhibitory neurons, and regulation of cell motility and cellular component movement in microglia (Fig. 3F).

### Regulatory network motifs and regulatory logics

The hierarchy of our cell-type GRNs suggests that the master regulators (top TFs) regulate crucial brain-related biological functions by regulating downstream TFs, which is a highly coordinated process. For instance, various network motifs have been found in GRNs, showing such a coordination pattern in which multiple TFs co-regulate target genes. To explore the extent of coordinated TF activities in our cell type GRNs, we computed the level of over or under-representation of all possible three-node network motifs (a triplet consisting of two TFs co-regulating a target gene, previously found to be enriched in many GRNs).

As depicted in Figure 4A, our brain cell type GRNs in AD and control broadly differ in their motif composition, and AD affects some of this composition (Data S3). For example, triplets in which two TFs target the same gene is over-represented in the microglial AD network relative to the control counterpart. On the other hand, enrichment of the motif in which two TFs are co-regulated by the same TF appears to be over-represented in microglia but underrepresented in oligodendrocytes (Fig. 4A), suggesting a possible disparity of TF-TF coordination across cell types. We were particularly interested in the feed-forward loops (FFL; TF1→TF2→TF3←TF1), as they have been often found to be biologically relevant in gene regulatory networks^19^. We found that FFLs are most conspicuous in the excitatory neurons and oligodendrocytes, but weakly enriched in inhibitory neurons and microglia (Fig. 4A). Interestingly, a zinc-finger transcription factor specificity protein 2 (SP2) is frequently found in FFLs (Fig. 4B). SP2 has been identified as a neural development gene^47^, but its role in AD has not yet been elucidated. Other TFs frequently found in FFLs across most cell type GRNs include FOXP1, RFX3, ZBTB18, and PPARA (Fig. 4B).

**Figure 4.**
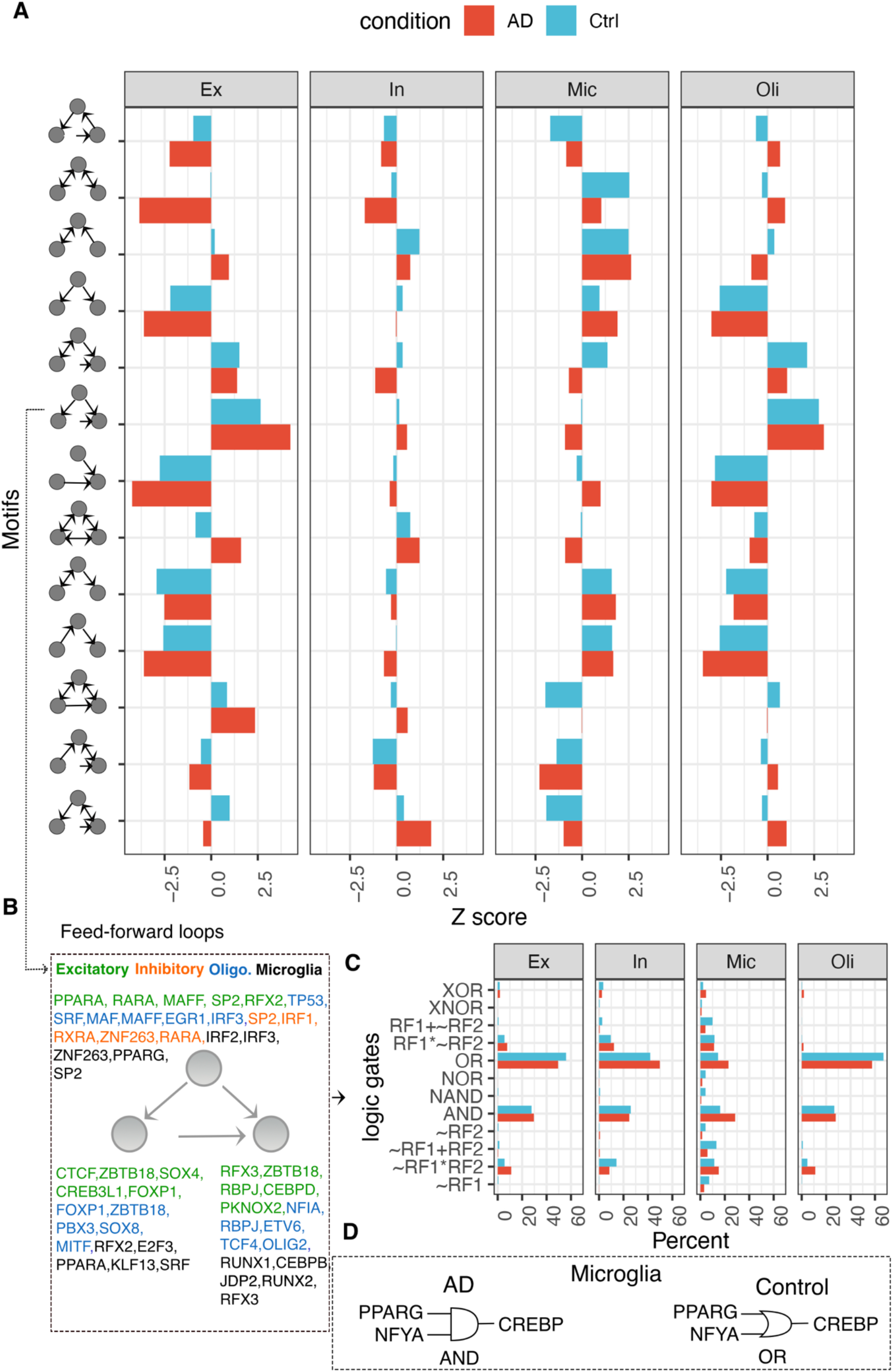
Network motifs and regulatory logic across cell types in AD. A) Barplots showing the enrichment (x-axis; Z-score estimates from random networks) of various three-node triplets (y-axis) in AD (red) and control (blue) conditions across all four cell types. B) Genes that frequently occur in feed-forward loops in cell type AD GRNs are depicted and colored uniquely for each cell type. C) Barplot showing the frequency (x-axis) of various logic gates (y-axis) active within the feed-forward loops in AD (red) and control (blue) conditions across all four cell type networks. D) Logic gate diagram showing PPARG-NFYA-CREBP triplet’s AND logic in AD and OR logic in control networks of microglia.

In addition to network motifs, we also investigated the cooperative logics of TFs that further reveal the TF-TF coordination mechanistically (beyond network structures like motifs). To this end, we applied our previous approach, Loregic^22^, to represent gene expression relationships in FFLs using logic gate models. In particular, logic gates describe gene regulation as a two-input one-output logical process^48,49^, where the expression level of regulatory factors (RFs) such as TFs are inputs and expression of target gene is the output. The logic gate has been a useful framework for studying cooperativity among RFs in human cancers, yeast and *E.* coli^19^. Thus, it would be interesting to investigate the logics behind TF cooperation in FFLs we discovered in our cell type GRNs. Fig. 4C shows that AND (high target expression only when both RF1 and RF2 are high) and OR (high target expression only when either RF1 and RF2 are high) represent more than 80% of all logics across all cell types, except in microglia. Microglia has more diverse logics than other cell types, many of which involve cooperative logics (i.e., RF1 and RF2 must be particular values to activate/repress target gene). For example, compared to other cell types, a larger fraction of logics in microglia involve RF1+~RF2, which means that target expression is low only when RF1 is low and RF2 is high is (see Table S1 in^22^ for explanation of these logics). An example of cooperative logics is the FFL consisting of PPARG-NFYA-CREBP which switches from uncooperative (OR) in control to cooperative (AND) in AD in the microglia network. In other words, PPARG and NFYA could be both required to active CREBP in AD, whereas either PPARG or NFYA can activate CREBP in healthy controls (Fig. 4D). PPARG is a ligand-activated nuclear receptor that coordinates lipid, glucose and energy metabolism and is upregulated in AD^50^. A GWAS study suggests that NFYA gene associates with late-onset AD^51^, and the CREBP gene functions in synaptic plasticity and memory formation and has been previously implicated in AD^52,53^. Thus, our logic analysis can further decipher the disease mechanisms of gene regulatory coordination of AD genes.

### Coregulated modules reveal cell type-specific drug-repurposed targets and gene functions in AD

Our analysis revealed features of the regulatory hierarchy and patterns of coordinated TF action in cell type GRNs, and changes in AD. It is important to also investigate non-TF genes, as they represent the larger component of the transcriptome. These genes lie at the bottom-most layer of the regulatory hierarchy as they have no outgoing links. We reasoned that interrogating this highly regulated core of target genes could illuminate dysregulated AD pathways and provide a handle on network modules.

We transformed the directed GRNs into undirected networks by connecting target genes that show high levels of ‘coregulation’ (estimated by calculating the overlap between the predicted regulators of every pair of target genes; see Methods). Using these networks of coregulated target genes, we tested the extent to which AD disrupts functional links between genes. We calculated the density (i.e., the ratio of observed to expected links) of the subnetworks induced by genes within carefully selected non-redundant GO BP terms. Then, comparing the densities of each GO BP term in control and AD networks allowed us to quantify the level of gain or-loss of ‘cohesiveness’ (i.e., interactions between GO BP genes became stronger or weaker in AD). This analysis highlighted several BP terms that significantly changed (permutation-based *p*-value < 0.001) densities across all cell types, with most in microglia (Fig. 5A). For example, in microglia, interactions between genes annotated to protein-membrane transport, lipid phosphorylation, cell aging, and other sugar metabolism related terms became stronger. Whereas GO BP terms that lost cohesiveness include actin cytoskeleton organization, regulation of interleukin-2 production, and B cell proliferation, among others (Fig. S4). In oligodendrocytes, interactions between genes involved in protein complex assembly, cytoskeleton organization, and neuron apoptosis became stronger, whereas interactions between genes involved in the Notch signaling pathway and transport activity became weaker (Fig. S5). In inhibitory neurons, genes involved in segmentation, cell cycle, and acid transport lost cohesiveness (Fig. S6). Whereas in excitatory neurons, genes involved in the cell cycle and response to biotic stimulus gained cohesiveness, while genes involved in apoptosis and response to fibroblast growth factor lost cohesiveness (Fig. S7).

**Figure 5.**
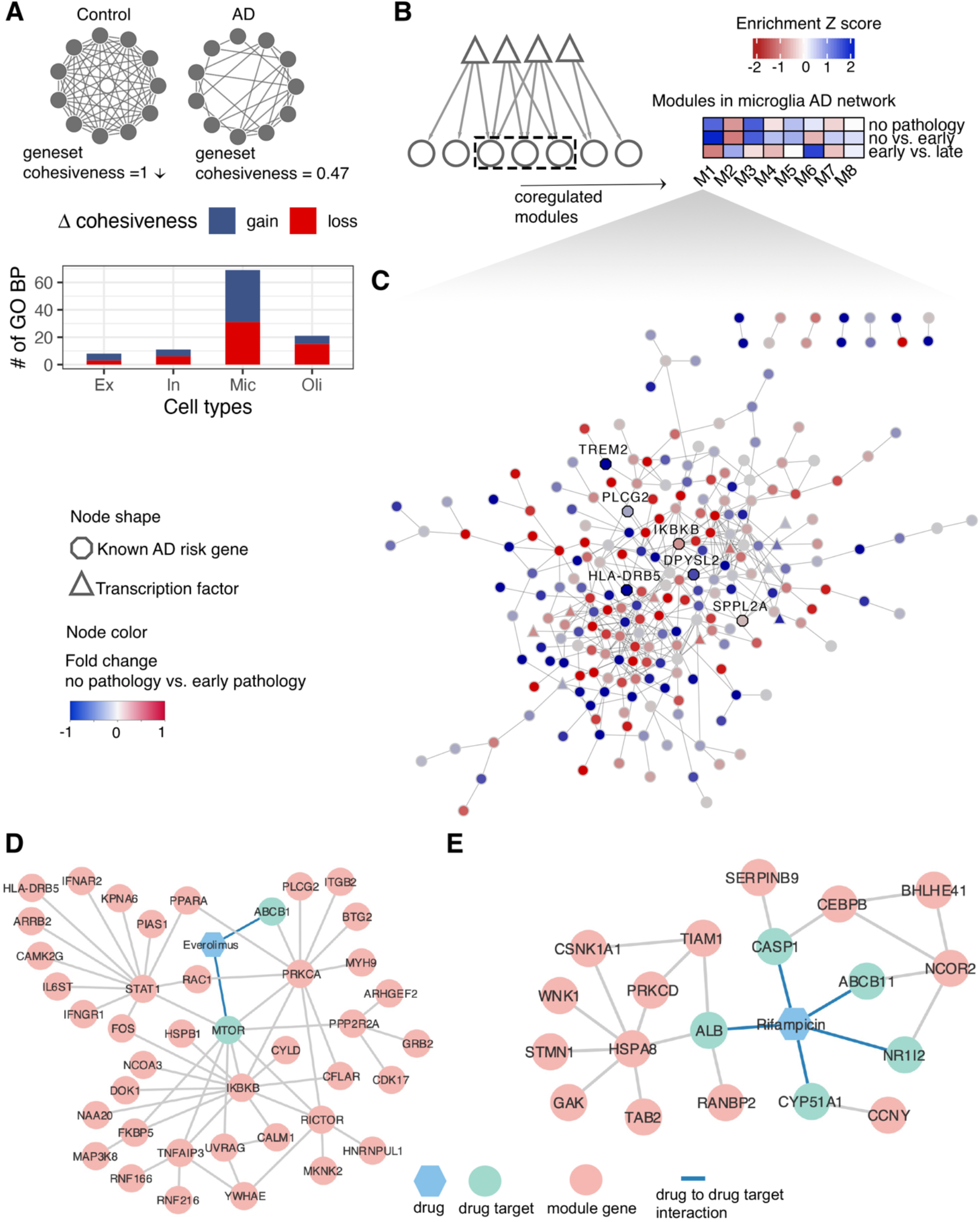
Coregulated gene modules reveal cell type-specific drug-repurposed targets and gene functions in AD. A) Illustration depicting the concept of gene set cohesiveness in a network. The bar plot below shows the number of gene ontology biological process terms (y-axis) that gain (blue) or lose (red) cohesiveness between control and AD networks across all cell types (x-axis; see Methods). B) A heatmap showing the enrichment of coregulated modules of the microglia AD network within differentially expressed genes in various AD pathologies. The average fold-change of genes within each module was transformed to a Z-score to derive the enrichment score. Negative and positive Z-scores indicate down- and up-regulation, respectively, of co-regulated modules (x-axis) in AD pathologies (y-axis). The grids of the heatmap are colored accordingly, with red indicating down-regulation and blue indicating up-regulation of the module. C) Visualization of genes in module 1 of the microglia AD coregulatory network. Each circle in the plot is a gene, with TFs depicted as triangles, known AD-genes in octagons, and other genes as ellipses. Nodes are colored according to fold change values in AD pathology (early versus no pathology) as shown in the key. D). Proposed mechanism-of-action for treatment of AD by everolimus using drug-target network analysis with microglia M1. E). Proposed mechanism-of-action for treatment of AD by Rifampcian using drug-target network analysis with microglia M4.

To explore the organization of target genes in cell type GRNs in more detail, we extracted network modules. We reasoned that the identification of modules will allow the use of modules rather than individual genes as units in our investigation of novel AD risk genes.

Rather than the typical approach of directly clustering gene expression data, we leveraged TF-target gene relationships embedded in our cell type GRNs to find functionally homogenous modules. Based on stringent evaluations of two clustering parameters (Fig. S8, S9, and S10; see Methods), we found on average 8 modules across all cell types, and these modules are significantly enriched (1% FDR) GO BP terms (Fig. S11A and S11B; Data S4 and Data S5). We also teased out AD modules as those that were significantly enriched (1% FDR) in disease ontology terms related to AD (Data S6). We found three AD modules each in excitatory and inhibitory neurons, and these modules are enriched in genes involved in processes related broadly to synaptic signalling, axonogenesis, and myelination (Data S5). The two AD modules in microglia are comprised of genes involved in the regulation of GTPase activity and various immune-related processes. However, we did not detect any AD module in oligodendrocytes. Nevertheless, our analysis shows that although many AD-risk genes functionally converge into common pathways with cell type specificity.

We next wanted to check the response of genes within these modules in three different AD pathologies; no pathology versus pathology, no pathology versus early pathology, and early pathology versus late pathology. We focused on modules in microglia for this analysis, as it showed a more extensive change in the cohesiveness of functional gene sets (Fig. 5A). We found that genes in microglia submodules 1 (M1) and 4 (M4) are more upregulated in the late stage of AD pathology compared with other microglia submodules (Fig. 5B). At the same time, the disease enrichment analyses (see Methods) demonstrated that M1 and M4 are significantly associated with AD (M1with q = 3.15E-02, M4 with q = 3.59E-02). M1 is enriched with genes involved in regulation of small GTPase mediated signal transduction and immune related processes according to the GO BP annotations. Interestingly, we found TREM2 as a part of M1, including other known AD-risk genes such as PLCG2, BIN1, IKBKB, DPYSL2, SPPL2A, and HLA-DRB5 (Fig. 5C). Triggering receptor expressed on myeloid cells 2 (TREM2) is a type I transmembrane protein expressed on the surface of microglia, binds to phospholipids^54^ and is hypothesized to be triggering the phagocytosis of Aβ plaques^55^. A recent study showed that TREM2 deficiency results in inhibition of FAK and Rac1/Cdc42-GTPase signaling critical for microglial migration^56^, testifying to the validity of M1. Neuroinflammation was proposed as one of the main mechanisms that were tightly associated with AD development^57^. KEGG pathway enrichment analysis showed that M1 was enriched with 12 immune pathways, including Fc gamma R-mediated phagocytosis, natural killer cell mediated cytotoxicity, toll-like receptor signaling pathway (Fig. S12A). Fc gamma R-mediated phagocytosis has been shown to play a role in ß-amyloid dependent AD pathology^58^. Toll-like receptor 4 (TLR4) activation was previously found positively correlated with the amount of accumulated ß-amyloid^59^. Furthermore, we found module M4 (Fig. S13) to be enriched with genes related to immune processes, such as response to chemokines, regulation of T cell migration, cytokine regulation (Fig. S12B)

Given the valid biological link of M1 and M4 to AD pathology, we next decided to predict drug candidates based on AD-related microglia submodules M1 and M4. With the well-defined network proximity approach^60^, we identified 170 and 34 candidate drugs with z_score < −2 and q < 0.05 from the total 2,891 U.S. FDA-approved or clinically investigational drugs (see Methods; Data S7). Interestingly, one of the drugs that show significant enrichment of its targets in M1 is Donepezil (*q* value 0.008), an approved AD drug that reversibly inhibits the acetylcholinesterase enzyme. Given that the Rho GTPase activity regulates the formation of Aβ peptides during disease progression^61^, our analysis raises an interesting hypothesis that the effect of Donepezil in improving the cognitive and behavioral signs and symptoms of AD might be executed via regulating GTPase signaling. Sildenafil, another top predicted drugs from M1, has recently been demonstrated as one promising treatment options that showed 69% reduction in developing AD after analysing MarketScan Medicare supplemental database which included 7.23 million individuals^31^. Everolimus, one mTOR inhibitor, was another top predicted drug from M1. Everolimus was discovered to bring down both human Aß and tau levels in the mouse model study^62^. Module M1 suggested that Everolimus’s target MTOR was directly connected with multiple key AD pathology regulators, such as inhibitor of nuclear factor kappa B kinase subunit beta (IKBKB), FKBP prolyl isomerase 5 (FKBP5) (Fig. 5D). One study in AD mouse model concluded that inhibiting IBKBK could help ameliorate activation of inflammatory and thus rescued cognitive dysfunction^63^. Level of FKBP5 was found to be positively correlated with AD development and FKBP5’s interaction with Hsp90 accelerated tau aggregation^64^. The same study also observed that decreased amount of tau in FKBP5^-/-^ mice. Rifampcian, one antibiotic drug, was top recommended according to module M4. One study found that Rifampcian was favourable for halting AD based on observations from both Aß and tau mouse models^65^. According to our protein-protein interaction network (see Method), similarly, multiple targets of Rifampcian were the direct neighbours of multiple proteins involved in AD development (Fig. 5E). CCCAT enhancer binding protein beta (CEBPB) was reported to modulate APOE’s gene expression and regulated APOE4 which was one major genetic risk factor for AD in one mouse model study^66^. Protein kinase C delta (PRKCD) which was one key protein in Fc gamma receptor-mediated phagocytosis pathways was found to regulate ß-amyloid dependent AD pathology^67^.

### Network-based machine learning prioritizes cell-type AD-risk genes and predicts clinical phenotypes

Our analysis shows several similarities and differences in cell type GRN structures patterns across cell types and between control and AD conditions. Decomposing the GRNs into individual components using standard network analysis metrics of centrality, hierarchy and modularity outlined key genes that potentially drive changes in cell type GRNs that underpin transcriptional phenotypes of AD. However, we were still lacking a uniform scoring to rank genes according to their potential association to AD using our cell type GRNs. To facilitate this, we leveraged known AD genes in the literature and asked if the regulatory patterns that characterize these could be learned. We reasoned that our GRNs are essentially high-level features extracted by integrating single-cell multi-omics data. Thus, regulatory patterns in these GRNs can be used to train machine learning (ML) algorithms. For example, this technique of using inferred network relationships as features for a learning algorithm has helped prioritize autism and hypertension genes in humans^68,69^ and stress-response TFs in plants^70^.

We used the random forest algorithm to train models that learned to discriminate between known AD genes and genes unrelated to AD using their interaction patterns with TFs as features (see Methods). We wanted to compare the accuracies in predicting known AD-risk genes across cell types and between control and AD networks. The distribution of balanced accuracies in 10 independent five-fold cross-validation tests indicates that the microglia AD network most accurately predicted known AD genes compared to other networks (Fig. 6A). The difference in mean accuracy between the control and AD networks of microglia is also the largest (Fig. 6A). The average accuracies of these models range between 57% to 68%. We ranked genes in each cell type model based on their predicted probabilities of being associated with AD (Data S8). The average probability of genes that were declared as differentially expressed in control versus AD by the original authors of the dataset^2^ is also relatively greater in microglial GRNs compared to other cell types (Fig. S14). Genes within the top 20% of the rankings in microglia AD network are involved in immune-related processes and hemopoietic functions, cell development, and lipid metabolism (Fig. 6B).

**Figure 6.**
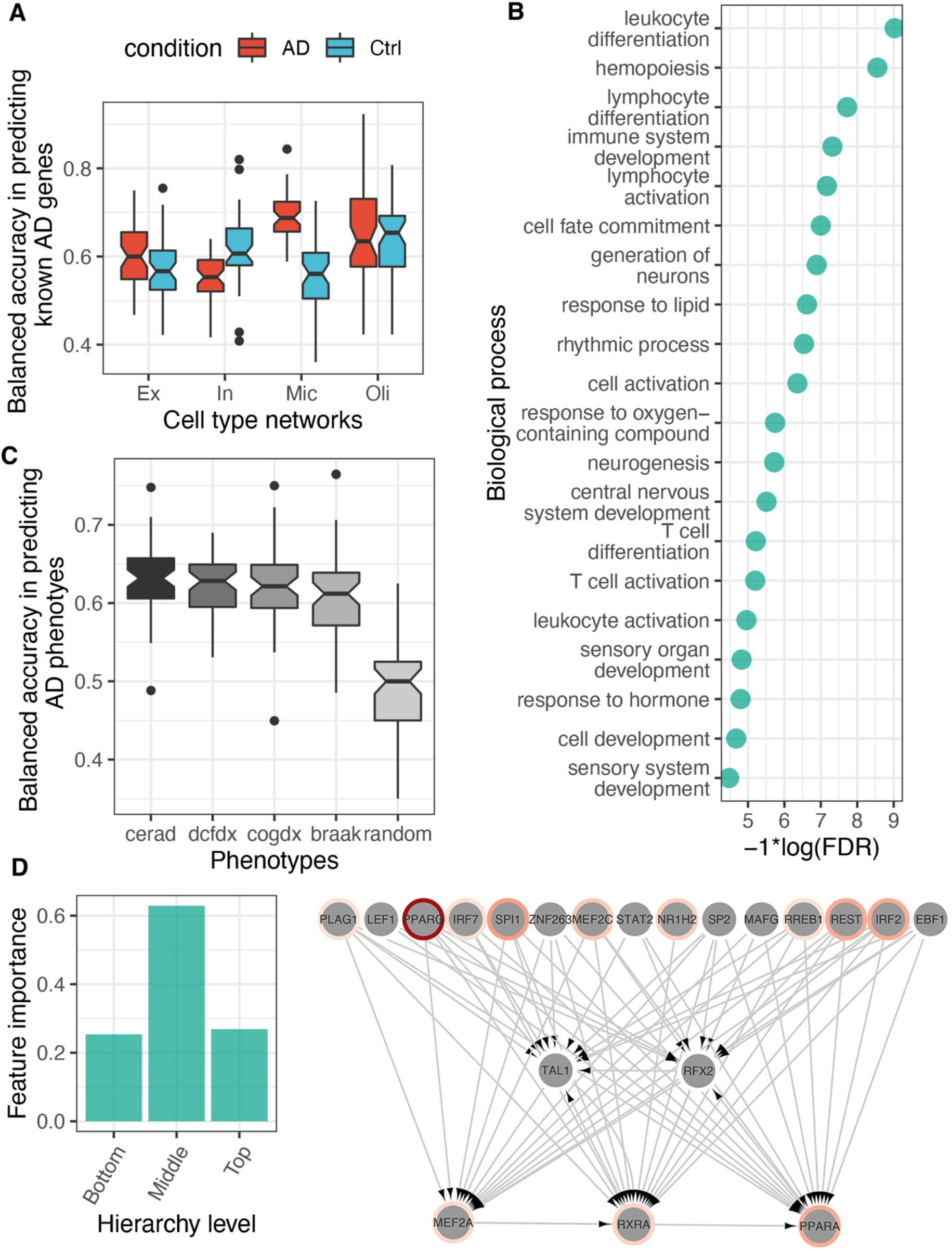
AD Gene prioritization and clinical phenotype prediction using networkbased machine learning. A) Boxplots showing the distribution of balanced accuracies (y-axis; obtained from 10 independent runs of five-fold cross-validation) in predicting known AD genes using interaction patterns in cell type GRNs as features (x-axis). B) Gene ontology biological process terms enriched within the top 20% predictions in the microglia AD machine learning (ML) model. The terms are depicted along the y-axis, and the FDR corrected *p*-values are shown along the x-axis. C) Genes were sorted according to their probability of being associated with AD in the microglia AD ML model, and the top 5% of the sorted list was used as features to predict AD phenotypes in an independent dataset (ROSMAP). The boxplots show the distribution of balanced accuracies (y-axis) obtained from these for four AD phenotypes (see Methods) and a set of randomly selected samples (x-axis). D) Average feature importance scores of TFs at the three hierarchy levels in the microglia AD network. E) Visualization of the subnetwork connecting top 10% TFs with highest feature importance scores in microglia AD network. Each grey node depicts a TF with border color set along a red gradient according to the disease-gene association score given in the DisGeneNet database (based on preliminary evidence collected from independent studies).

To evaluate these rankings more stringently, we asked if the expression levels of the topranked genes could be used to predict clinical phenotypes of AD. We utilized RNA-seq data from the ROSMAP cohort^71^ to predict AD phenotypes, including Braak stages that measure the severity of neurofibrillary tangle (NFT) pathology, CERAD scores that measure neuritic plaques, diagnosis of cognitive status (DCFDX), and cognitive status at the time of death (COGDX). Figure 6C shows the distributions of accuracy scores in predicting these phenotypes using top 5% genes in our rankings from the microglial AD model as features. Our model shows that these genes can classify AD phenotypes with more than 60% accuracy, larger than the model built using randomly selected genes (Fig. 6C).

By analyzing TFs separately, we observed that those in the middle layer of the regulatory hierarchy record the highest feature importance scores (served as the best predictors in the model; Fig. 6D), indicating their prominent role in regulation of gene expression in AD (Data S9). Visualization of the subnetwork among these TFs revealed known AD genes and interaction patterns (Fig. 6E). For example, we found two genes from the peroxisome proliferator-activated receptors (PPARG and PPARA) in this subnetwork. PPARs function in inflammation and immunity^72^, coordinate glucose and energy metabolism^73,74^, and are known to positively influence AD pathology. In addition to this, PPARA regulates genes involved in fatty acid metabolism and activates hepatic autophagy^75^. Other interesting TFs in this subnetwork include SPI1, a well-known TFs involved in microglial development and activation^76^ and has been implicated in AD in GWAS^77^. Interestingly, our analysis prioritized several TFs with no previous direct associations to AD in databases. Some such examples include TAL1, RFX2, LEF1, SP2, STAT2, ZNF263, MAFG, and EBF1. Therefore, it would be interesting to further investigate the link between these TFs and AD.

## Discussion

We developed an integrative analysis framework on single-cell network biology and applied it to cell type gene regulatory networks (GRNs) in AD (Fig. 1). The networks were predicted by single-cell multi-omics data, including scRNA-seq, cell type chromatin interactions, and TF binding sites on open chromatin regions. We identified the AD-specific network characteristics changes such as hub genes and regulators, modules, and hierarchies, suggesting their vital roles for cellular and molecular functions in AD pathogenesis by analyzing those networks. Further, we revealed that using those cell type networks also improved the prediction of potential novel AD genes, clinical phenotypes, and drug targets.

Our analysis of gene centrality metrics suggests that different cell types employ a different regulatory apparatus governed by ‘master regulators’ that contributes to cell viability, and therefore overall brain fitness by regulating the expression of many target genes. This apparatus does not seem to be perturbed by the occurrence of AD, which makes sense considering previous studies on gene essentiality and lethality. Interestingly, distinct TFs with high betweenness centralities seem to modulate cell type-specific signals. Furthermore, non-TF genes, or in-degree centralities exhibit the most prominent differences between AD and control brains, indicating that these genes may contribute to pathway-level changes in AD progression. Enrichment of critical brain-specific biological processes, such as synapse organization and immune-related processes, within these genes also reflects characteristics of AD pathology. Overall, centrality analysis can delineate ‘master regulators’ that are potentially involved in maintaining biological processes essential for cell type function in healthy and AD individuals.

Our analysis shows that the brain GRNs are hierarchical in structure and uncovered the hierarchy height of brain TFs for the first time. Our analysis suggests that the levels on which TFs operate are generally robust to AD, and subtle changes in the expression of TFs at the top and bottom levels seem to modulate cellular signals underlying typically observed AD phenotypes. GO BP enrichment analysis also suggests that TFs in the bottom layer are involved in relevant processes like ‘synapse organization’, ‘neuron projection development’ and ‘axon development’ in neuronal cell types, ‘neutrophil immunity’, ‘endocytosis’, ‘cellcell adhesion’ and ‘actin organization’ in microglia, and ‘neuron projection development’, ‘cell morphogenesis’ and ‘axonogenesis’ in oligodendrocytes. However, the middle-level TFs seem to be most active and perhaps cooperate and coordinate with other TFs to target a relatively larger number of genes. We noted that hierarchy height for TFs estimated using ChIP-seq datasets better reflected a tri-modal distribution^43^. In our analysis, we found fewer TFs in the middle layer and many TFs at the top layer. This could perhaps be due to many indirect TF-TG correlations that naturally arise in expression data. Another strategy to infer regulatory hierarchies more accurately would be to apply a simulated annealing procedure to the full network (including non-TF genes) to get better estimates on the actual number of hierarchies in a given network^78^. Such an analysis requires a considerable amount of computational runtime beyond our timeframe, given that we analyzed a total of eight networks. Nevertheless, our analysis’s distribution of TF hierarchies is statistically significant compared to random networks. Moreover, whether the occurrence of fewer TFs in the middle levels is a feature of single-cell GRNs or just noise due to indirect correlations could only be evaluated based on new data from cell type-specific TF-DNA binding data.

Our TF-centric analysis indicates extensive dysregulation in the microglia network. For example, microglia has a unique set of TFs with high betweenness centrality (Fig. 2A) and the largest rewiring between control and AD networks (Fig. 3C). Therefore, we wanted to investigate dysregulation at the level of non-TF genes, as these genes represent the core brain pathways. Indeed, using coregulation levels as a proxy for functional relatedness, we confirmed that biological processes are most dysregulated in microglia networks (Fig. 5A). This also testifies that our approach of utilizing known functional gene sets (e.g., GO terms) as biologically coherent components and using subnetwork density as a metric to gauge gene set activity is an excellent approach to highlight individual cell types. We found that the cell type networks are highly modular, and the organization of modules is largely distinct, as the overlaps between cell type modules were minimal (Sup Fig.). We chose to investigate microglial module 2 further as this was the only module explicitly upregulated in the early stages of AD and also statistically enriched with genes that support known AD biology. Lipid metabolism has been previously implicated in AD^79^, and the role of microglia in lipid metabolism is also previously suggested^80^. Thus, we anticipate that microglial module 2 genes in our analysis could potentially have pharmacologic applications for early intervention and drug research to target lipid dysregulation.

The cell type GRNs we inferred in this study, together with extensive prior genetic knowledge on AD, presented us with a unique opportunity to identify patterns of regulatory interactions that characterize AD. Inspired by previous network-based machine learning approaches, we developed an approach that leverages regulatory interactions of known AD genes as the ground truth to find other similar yet uncharacterized AD genes. Our approach correctly prioritized microglial genes related to lipid metabolism and hemopoietic function; these are well-known biological processes disrupted in AD. However, the average accuracy of our best model (~0.68) is lower than what is typically expected from such models. This low accuracy could have arisen because we utilized network data from a single-cell type to train the models, effectively neglecting the possible functional role of other cell types in AD^81^. Unfortunately, our models do not capture this complexity even when different neuronal and glial networks were integrated into a single prediction model (data not shown). This could be due to the fact that our cell type networks lack chromatin interaction data in AD. Nevertheless, we are hopeful for the future as more single-cell multi-omics data accumulate in the context of AD, enabling us to further refine our network models.

Overall, our integrated single-cell network analysis approach identified key genes and cellular themes that corroborate many aspects of AD biology. This shows that gene regulatory networks extracted from single-cell data can reveal molecular systems often hidden in gene networks derived from bulk datasets. For example, our networks revealed extensive network rewiring disrupting key biological processes mainly in microglia. As such, our approach can pinpoint cell type-specific genes that could potentially play a key role in governing disease-induced changes of pathways (e.g., lipid metabolism). As the single-cell technology further advances our ability to capture multi-modal genomic data with unprecedented precision, we anticipate that network biology applied to such single-cell functional genomics data will enhance precision medicine. Single-cell sequencing assays offer solutions to two main requisites for statistical inference of reliable gene networks; large sample size and context-specificity (unifying biological theme defined by the underlying datasets). While bulk RNA-seq datasets could provide researchers with a large enough sample size, the context-specificity is often ambiguous in publicly available datasets^82^. Single-cell technology, by design, generates volumes of data from each individual in the study. As such, pooling cell type samples from individuals is currently recommended by not required for cell type network inference. Thus, patient-specific gene networks could be possible in the coming years, which will enable us to predict a clinical outcome better (e.g. drug response) based on network activity of target components (e.g. drug targets)^3^. Furthermore, since we already collect patient-specific data from other modalities (e.g. imaging, behavioral and clinical), fusing genetic network models with models from non-genomic modalities could resolve overlapping disease features better. Our network biology approach provides a method to investigate disease genes from single-cell data and lends itself to be used as a template for genomic feature engineering for advanced AI-based integrative models.

## Methods and Materials

### Single-cell datasets and data processing

We obtained previously published single-cell gene expression data for major cell types including excitatory and inhibitory neurons, microglia, and oligodendrocyte from individuals with Alzheimer’s disease pathology and healthy controls^2^. Precisely, the dataset consists of single-nucleus RNA-sequencing (snRNA-seq) of samples from the prefrontal cortex of 24 individuals with varying degrees of AD pathology and 24 age-matched controls with no AD pathology. We removed genes that were expressed in less than 100 cells and normalized the data using Seurat 4.0^83^. We then applied MAGIC^84^ to address dropout events by imputing the missing gene expression values and filtered lowly expressed genes to create cell type gene expression matrices. In addition, we also obtained other omics data, including cell type chromatin interaction maps^32^, transcription factor binding sites^33^, and cell type open chromatin regions^85^.

### Gene regulatory network inference for brain cell types from multi-omics

We sought to integrate single-cell transcriptomic, chromatin interaction, TF binding sites, and open-chromatin regions to predict directed edges from transcription factors (TFs) to target genes (TGs). We used our scGRNom (single-cell gene regulatory network prediction from multi-omics) pipeline to perform this integration^86^. First, the scGRNom function *scGRNom_interaction* was supplied with cell type chromatin interaction data to predict all possible interactions between enhancers and promoters. Then, reference networks for each cell type were obtained by locating human TF binding sites (TFBS) within the identified interacting regions using the function *scGRNom_getTF*. Subsequently, the reference networks along with the single-cell gene expression matrix were supplied to the *scGRNom_getNt* function to predict TF-target genes for each cell type. The *scGRNom_getNt* uses elastic net regression to infer TF-target gene relationships. To identify the most confident edges, we filtered the target genes with mean squared error > 0.1 and absolute elastic net coefficient < 0.01.

### Analysis of cell type GRN characteristics

#### Centrality analysis

Three measures of network centrality were used to gauge the importance of genes in each network. The indegree and outdegree of genes in a given network were calculated as the number of incoming TFs for each target gene and the number of target genes for each TF, respectively. The betweenness centrality was calculated by counting the number of times a given gene appears within the shortest paths of two other genes in a given network. The centrality scores for each network were scaled between 0 and 1 to make the scores comparable across cell types. To calculate fold change in centrality scores in AD versus control network of each cell type, we first replaced missing values (genes found in AD network but not in control or vice-versa) with the number that equals 1% of the smallest observed centrality score in both the networks to avoid dividing by 0. The fold change of a given gene was then calculated as the binary logarithm of the gene’s normalized centrality score in the AD network divided by the control network. Genes with absolute scores > 0.5 were used for functional enrichment analysis (described below). All networks were treated as directed and the igraph R library was used to estimate gene centrality scores.

#### Hierarchy analysis

We used the hierarchy height (*h; outdegree - indegree*) of TFs to probe the direction of information flow in each network. The following analysis was performed on only TF-TF networks (TG is also a TF). The normalized *h* metric was calculated as^43^

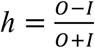

Where *O* = outdegree and *I* = in-degree of a TF. With this metric, TFs with *h* between 1 and 0.33 were classified as the top-level regulators, TFs with *h* between 0.33 and −0.33 were classified as the middle-level regulators, and TFs with *h* between −0.33 and −1 were classified as bottom-level regulators. The significance of the distribution of *h* metric of TFs, which was trimodal across most cell types, was calculated from the distribution of *h* in 1000 random networks (KS tests). The random networks were generated by preserving the observed edge density in each network.

#### TF rewiring analysis

To quantify the difference between sets of predicted targets of a TF in control versus AD networks, we calculated the rewiring score as^43^

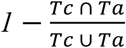

Thus, a high rewiring score of a TF means that its targets in the control network (*Tc*) and AD network (*Ta*) exhibit little overlap.

### Network motifs and regulatory logic analysis

Motif analysis was used to identify specific interaction patterns in the networks. We focused on subgraphs containing three genes, which were collated into 13 isomorphic classes. The number of times each class occurred in each network was recorded using the mfinder tool ^87^. The Z score of the distribution was estimated from 1000 random networks. Due to a large number of networks, we set the sampling parameter to 100 to obtain a fast approximate motif analysis of the networks. To characterize TF more of action in feed-forward loops, we applied logic circuit models using the Loregic algorithm^22^. Loregic classifies TFs into regulatory triplets (two TFs and a target gene forming an FFL in our analysis) and identifies the logic gate model (e.g. AND, OR, XOR etc.) most consistent with the cross-sample expression of each triplet. Loregic requires a binarized form of expression data as input to score the logic gate models. For each cell type gene expression matrix, we selected 100 cells with the highest variance as inputs to Loregic. 100 cells were selected to account for the uneven distribution of different cell types in the full expression matrix. This also allowed us to reduce the overall runtime of the Loregic algorithm. Loregic outputs a gate consistency score for each of the 16 possible logic gate models. For each triplet, we selected the gate model with the highest consistency score as the gate consistent for the triplet. Gates with ties in the consistency score were regarded as gate inconsistent. The statistical significance of consistent gates was estimated by replacing the target gene in each triplet with a random gene from the corresponding network and calculating the fraction of time the gate consistency score of the randomized triplet was greater than or equal to the empirical score. Consistent gates with *P* <=0.01 were reported.

### Calculation of gene-set cohesiveness and identification of coregulated gene modules

Our network dataset contained directed networks in which TFs are one set of nodes with outgoing links and target genes as another set of nodes with incoming links. Because TFs can also have incoming links, the networks we had at hand were essentially structured as mixed bipartite graphs. We transformed these directed graphs into undirected graphs by connecting target gene pairs if they had a considerable overlap between their predicted regulators. The overlap between the predicted regulators of a given gene pair was estimated using the Jaccard’s Index (JI) and set the edge-weight. Using these weighted graphs, the gain or loss of cohesiveness within functional gene sets (GO BP terms) was estimated as follows. First, for a given gene set, a subnetwork depicting edges within the gene set was extracted. Then, the normalized network density of the subnetwork was calculated as the sum of edge weight divided by the total genes in the gene set. These operations were performed across control and AD networks of all cell types. Finally, change in gene set cohesiveness was calculated as the log ratio of density in the AD network divided by density in the control network. The statistical significance of Δ cohesiveness was calculated by randomly sampling the gene set from the background of all genes in the AD networks and calculating the picking genes from all gene sets with a fold-change greater than 0.5 were reported in Figure 5A.

Then, the adjacency matrix holding target genes in rows and columns and JI values in the cells was supplied to the WGCNA algorithm to detect coregulated gene modules^88^. The detection of reliable modules will depend on two critical parameters: the edge-weight threshold (EWT) to maintain high scoring edges and filter noise arising due to indirect regulations and the minimum module size (MMS) parameter. We wanted the MMS to be large enough (atleast 10 genes) to objectively test the functional relevance of resulting modules using statistical enrichment but not too large to include bifurcated components of large metabolic pathways into the same modules. Therefore, we tested a range of EWT (between 0.1 and 0.9) and MMS values (between 10 to 100) for every cell type network to obtain the best possible solution. We asked what combination of EWT and MMS detects the largest number of functionally relevant gene modules while retaining as many original genes as possible to avoid information loss. The functional relevance was tested by counting the fraction of detected modules that could be annotated using statistical enrichment of GO BP terms. Based on these evaluations, we found an EWT of 0.2 (20% overlap between predicted regulators of a TG-pair) and an MMS of 30 yields the best network clustering solution (Fig. S4, S5, and S6. The *blockwise module function* of WGCNA was invoked with ‘agglomerative clustering using average linkage’ as the clustering algorithm.

### Functional and disease gene enrichment analysis

The human gene ontology biological process (GO BP) annotations^89^, propagated along ‘is_a’ and ‘part_of’ relationships were obtained^90^. Enrichment of query genes (e.g. top central genes, module genes etc.) within a given functional geneset (GO BP term or modules) was calculated using hypergeometric tests, using all genes present in the corresponding network as the background. The resulting p-values were corrected for multiple testing using the Benjamini-Hochberg method^91^. Note that for GO, apart from propagating parent-child relationships, we also removed geneset terms that annotate more than 500 and less than 10 genes for enrichment analysis.

### Network proximity for drug prediction

We assembled drugs from the DrugBank database relating to 2,891 compounds^92^. To predict drugs with interested modules, we adopted the closest-based network proximity measure^60^ as below

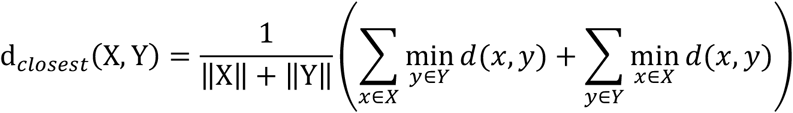

where d(x,y) is the shortest path length between gene x and y from gene sets X and Y, respectively. In our work, X denotes the interested modules, Y denotes the drug targets (gene set) for each compound. To evaluate whether such proximity was significant, the computed network proximity is transferred into z score form as shown below:

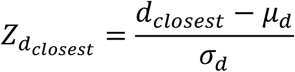

Here, *μ_d_* and *σ_d_* are the mean and standard deviation of permutation test with 1,000 random experiments. In each random experiment, two random subnetworks *X_r_* and *Y_r_* are constructed with the same numbers of nodes and degree distribution as the given 2 subnetworks X and Y separately, in the protein-protein interaction network.

### Protein-protein interactome (PPI) network

To build the comprehensive human interactome from the most contemporary data available, we assembled 18 commonly used PPI databases with experimental evidence and the in-house systematic human PPI that we have previously utilized: (i) binary PPIs tested by high-throughput yeast-two-hybrid (Y2H) system^93^; (ii) kinase-substrate interactions by literature-derived low-throughput and high-throughput experiments from KinomeNetworkX^94^, Human Protein Resource Database (HPRD)^95^, PhosphoNetworks^96^, PhosphositePlus^97^, DbPTM 3.0 and Phospho.ELM^98^; (iii) signaling networks by literature-derived low-throughput experiments from the SignaLink2.0^99^; (iv) binary PPIs from three-dimensional protein structures from Instruct^100^; (v) protein complexes data (~56,000 candidate interactions) identified by a robust affinity purification-mass spectrometry collected from BioPlex V2.0^101^; and (vi) carefully literature-curated PPIs identified by affinity purification followed by mass spectrometry from BioGRID^102^, PINA^103^, HPRD^104^, MINT^105^, IntAct^106^, and InnateDB^107^. Herein, the human interactome constructed in this way includes 351,444 PPIs connecting 17,706 unique human proteins.

### Machine-learning model for AD-gene prioritization

We sought to utilize network connectivity patterns in cell type regulatory networks to make predictions on disease-gene associations. First, we downloaded known disease-gene associations listed in the DisGenNet database^108^ and extracted all genes linked with the keyword ‘Alzheimer’. The DisGenNet database ranks gene-disease associations using a metric that quantifies the level of evidence in published literature. 16% (3481 out of 21666) of all genes in the database are linked with AD, with gene-disease associations scores ranging from 0.01 (not strong evidence) to 0.9 (strong evidence). We selected AD genes with scores greater than 0.1 (top 20%) as positive examples to build the binary classifiers. Then, rather than randomly selecting negative samples, we further analyzed the DisGenNet database to identify genes that are likely not associated with AD. To do this, we calculated overlaps between diseases and selected genes strongly associated with diseases that have minimal overlaps with AD (disease-disease Jaccard’s overlap < 0.1). From this pool of ‘likely not AD-associated’ genes, we randomly selected negative examples equal to the number of positive examples to build classifiers not biased by class-size. Then, each GRN was transformed into a non-symmetrical adjacency matrix *A*, with TFs (*i*) in columns and TGs (*j*) in rows and the cell *Aij* containing the predicted edge score (absolute coefficient of elastic net regression from scGRNom) of the corresponding TF-TG pair. The subset of *A* with rows containing our positive and negative samples was extracted as the feature matrix, *F*. To include TFs that do not have any in-degrees (not regulated by other TFs in our networks) in *F*, we assigned an edge score equal to 1% of the minimum edge score in the corresponding network. This allowed us to label TFs with no upstream regulators and include them in prediction models. Then, using the vector of edge scores of each sample in *F* as the feature vector, we trained a random forest classifier to discriminate between positive and negative samples. The balanced accuracy of the model was tested using 10 independent runs of fivefold cross-validation. The average balanced accuracy (total 50 trials) was recorded and plotted. The predicted probability of class output from the classifier was used to rank all genes. The feature importance score was measured as the Gini impurity. The Gini impurity metric estimates the probability of classifying a sample incorrectly, and is calculated as

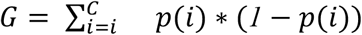

where *C* is the total number of classes (2 in our case) and *p*(*i*) is the probability of picking a sample in class *i*. The accuracy and *G* were recorded for each cell type GRN in both conditions. The most accurate cell type model was chosen as the one with the highest average accuracy (AD microglia network in our study) and used to predict the probability of AD association of the remaining unlabeled genes along with TFs with the largest feature importance scores.

### Prediction of clinical phenotypes

To predict AD phenotypes, we utilized the original RNA-seq data from the ROSMAP study (55,889 Ensembl gene ids for 640 post-mortem human samples) on an Alzheimer’s disease case-control cohort for the Dorsolateral Prefrontal Cortex (DLPFC) brain region. We obtained permission from ROSMAP to use this data (available on synapse.org (ID: syn3219045). We mapped the Ensembl genes ids to Entrez gene identifiers, averaged the gene expression values for Ensembl gene identifiers that mapped to the same Entrez identifiers, and removed unmapped Ensembl identifiers. Ultimately, we found 26,017 genes (with unique Entrez IDs). Only 638 out of 640 individual RNA-Seq samples mapped to population phenotypes. Our final DLPFC dataset thus contained gene expression values for 26,017 genes for 638 samples. Then, using normalized gene expression values of top 5% ranked genes from the microglia AD-gene classification model (described above) as the feature vectors, we trained random forest classifiers to predict various AD phenotypes. The following coding was used: cogdx (4 and 5 versus 1), braak (0,1,2 versus 5,6), and cerad (1 versus 3,4). The classifier accuracy was evaluated using 10 independent runs of five-fold cross-validations, as described above.

## Acknowledgements

This work was supported by National Institutes of Health grants, R01AG067025, R21CA237955, R03NS123969 and U01MH116492 to D.W., P50HD105353 to Waisman Center, and the start-up funding for D.W. from the Office of the Vice Chancellor for Research and Graduate Education at the University of Wisconsin-Madison.

## Data availability

All our results are provided in Supplementary Datasets. All processed data are available at https://github.com/daifengwanglab/scNET. All other data are available from the corresponding author on reasonable request.

## Code availability

The codes for our analyses are available at https://github.com/daifengwanglab/scNET.

## Contributions

D.W. conceived the study. D.W. and C.G. designed the experiments. C.G. performed the experiments and analyzed the data. J.X. and F.C. performed drug-target interaction analysis. C.G., J.X., S.K., F.C. and D.W. wrote the manuscript. T.J, S.K, X.L., and S.A. contributed to data analysis. All authors read and approved the manuscript.

## Supplementary information

Data S1: Gene centralities across all cell type GRNs

Data S2: Hierarchy of TFs across all cell type GRNs

Data S3: Results of motif enrichment of cell type GRNs

Data S4: Gene module memberships

Data S5: GO enrichment results of network modules

Data S6: Disease ontology enrichment of network modules

Data S7: Drug prediction results

Data S8: Predicted probabilities of network-based AD gene classifiers

Data S9: Feature importance scores of TFs in microglia AD network

